# Sennoside A differentially regulates mast cell activation and attenuates gut allergic inflammation

**DOI:** 10.64898/2026.09.23.753939

**Authors:** Fathima Nawaz, Cheyanne Woodrow, Ana C. Roginski, Esther B. Florsheim

## Abstract

**Introduction:** Mast cells are central effectors of IgE-mediated allergic responses and reside in barrier tissues where they encounter diverse environmental compounds. How naturally occurring bioactive compounds influence mast cell function under resting versus allergic conditions remains poorly defined.

**Methods:** We compared the effects of cholera toxin, aconitine, and sennoside A on viability, degranulation, and cytokine production in murine bone marrow-derived mast cell. We then examined the acute physiological effects of oral sennoside A and its effects in an ovalbumin-induced mouse model of food allergy.

**Results:** None of the compounds reduced mast cell viability at the concentrations tested. Sennoside A, but not cholera toxin or aconitine, induced modest degranulation in resting mast cells without increasing TNF-α or IL-6 secretion. In contrast, sennoside A markedly suppressed IgE-dependent antigen-induced degranulation, TNF-α and IL-6 secretion, and *Il1b* expression. Acute oral sennoside A induced dose-dependent diarrhea and hypothermia and increased colonic *Tnfa*, *Il1b*, *Il33*, and *Hdc* expression. In allergic mice, sennoside A attenuated allergen-induced hypothermia, reduced duodenal *Tpsab1* expression, and decreased circulating total IgE, OVA-specific IgE, and OVA-specific IgG1, while increasing *Il5* expression.

**Conclusion:** Sennoside A differentially regulates mast cell responses according to activation state and attenuates features of experimental food allergy. These findings identify a previously unrecognized immunomodulatory effect of sennoside A on IgE-driven mast cell activation. However, the mechanisms and the contribution of mast cells to the *in vivo* phenotype remain to be defined.

## INTRODUCTION

Innate immunity is central to the detection of and protection against pathogens, but how this system responds to non-pathogenic toxic compounds remains poorly understood. Toxins are ubiquitous in nature and are produced across all biological kingdoms, representing a persistent challenge to animal fitness. This challenge is particularly acute at barrier tissues, where essential activities such as feeding inevitably expose animals to potentially harmful chemicals. If accidentally ingested, noxious compounds must be rapidly neutralized or eliminated to limit tissue injury. Canonical xenobiotic defense pathways, including hepatic biotransformation and renal excretion, metabolize and clear potentially harmful substances (1,2). Immune responses may provide an additional layer of protection against noxious exposures (3,4). In particular, allergic immunity has been proposed to function as a biological food quality control system, mounting protective responses against food antigens associated with harmful or potentially toxic substances (5). Consistent with this idea, IgE- and mast cell-dependent sensing of ingested allergens promotes antigen-specific avoidance behavior, providing a mechanism to limit subsequent exposure to potentially harmful foods (6). Whether and how innate immune cells directly detect and respond to ingested toxic compounds, however, remains less clear. Defining these responses may reveal defense mechanisms that extend beyond classical recognition of pathogen-associated molecular patterns.

Plant secondary metabolites illustrate the biological complexity of these exposures (7). Many evolved as chemical defenses (8) and can be harmful at one dose while producing therapeutic or immunomodulatory effects under other conditions (9,10). Aconitine, an alkaloid from *Aconitum* species, disrupts voltage-gated sodium channel function and can cause severe neurological and cardiovascular toxicity (11,12), yet it has also been reported to alter inflammatory pathways (13). Sennoside A, a dianthrone glycoside found in *Senna* species and rhubarb, is best known for its laxative activity after ingestion. Sennoside A has also been reported to suppress inflammatory responses in experimental disease models, including reductions in TNF-α, IL-6, IL-1β, and MCP-1 during metabolic inflammation (14). These observations raise the broader question of how immune cells at barrier tissues respond to bioactive plant compounds.

Among innate immune cells, mast cells are particularly well positioned to respond rapidly to environmental and dietary compounds encountered at barrier tissues. Mast cells store preformed bioactive mediators that can be released within minutes of activation and are best known for their central role in IgE-mediated allergic responses (15,16). Recent work has further shown that intestinal mast cells are functionally specialized by the mucosal environment and mediate IgE–FcεRI-dependent anaphylactic responses to ingested antigens, with cysteinyl leukotrienes serving as key effector molecules (17). However, mast cells can also respond directly to noxious substances independently of IgE. Mast cell proteases contribute to protection against several animal venoms (18), and microbial toxins including *Staphylococcus aureus* δ-toxin, Group B *Streptococcus* pigment, *Clostridioides difficile* toxin TcdB, and streptolysin O can directly alter mast cell function (19–22). These observations raise the possibility that naturally encountered bioactive compounds may influence mast cell responses outside canonical allergen recognition and, importantly, may modify subsequent IgE-dependent activation.

In this study, we examined the effects of three biologically and chemically distinct compounds—cholera toxin, aconitine, and sennoside A—on mast cell viability and activation. We found that mast cell responses were compound-specific: cholera toxin and aconitine did not induce detectable degranulation under the conditions tested, whereas sennoside A induced degranulation in resting mast cells. Unexpectedly, sennoside A had the opposite effect during IgE-dependent activation, markedly suppressing antigen-induced mast cell degranulation and inflammatory mediator production. We therefore examined the physiological effects of oral sennoside A and its impact in an experimental model of food allergy. Sennoside A induced acute intestinal and systemic responses in otherwise healthy mice but attenuated allergen-induced hypothermia and reduced several immunological features of gut allergic inflammation. These findings identify sennoside A as a context-dependent modulator of mast cell function and reveal distinct effects of this plant-derived compound under basal and allergic inflammatory conditions.

## RESULTS

### Sennoside A selectively induces mast cell degranulation without reducing mast cell viability

To establish an *in vitro* system for testing mast cell responses to chemically distinct compounds, bone marrow cells from female BALB/c mice were cultured with IL-3 and stem cell factor (SCF) for six weeks (Supplementary Figure 1). Cultures used for experiments contained >90% CD117⁺FcεRI⁺ cells, consistent with a mature bone marrow–derived mast cell (BMMC) population.

We first asked whether sennoside A, aconitine, or cholera toxin compromised mast cell viability. BMMCs were exposed for 24 h to sennoside A (25 or 250 μg/mL), aconitine (50 or 100 μg/mL), or cholera toxin (0.5 or 2 μg/mL), followed by Zombie Yellow staining and flow cytometry (Fig. 1a). Boiled cells served as a nonviable positive control and were nearly uniformly Zombie Yellow-positive (Fig. 1b). None of the three compounds significantly reduced mast cell viability at the concentrations tested (Fig. 1b).

**Figure 1.**
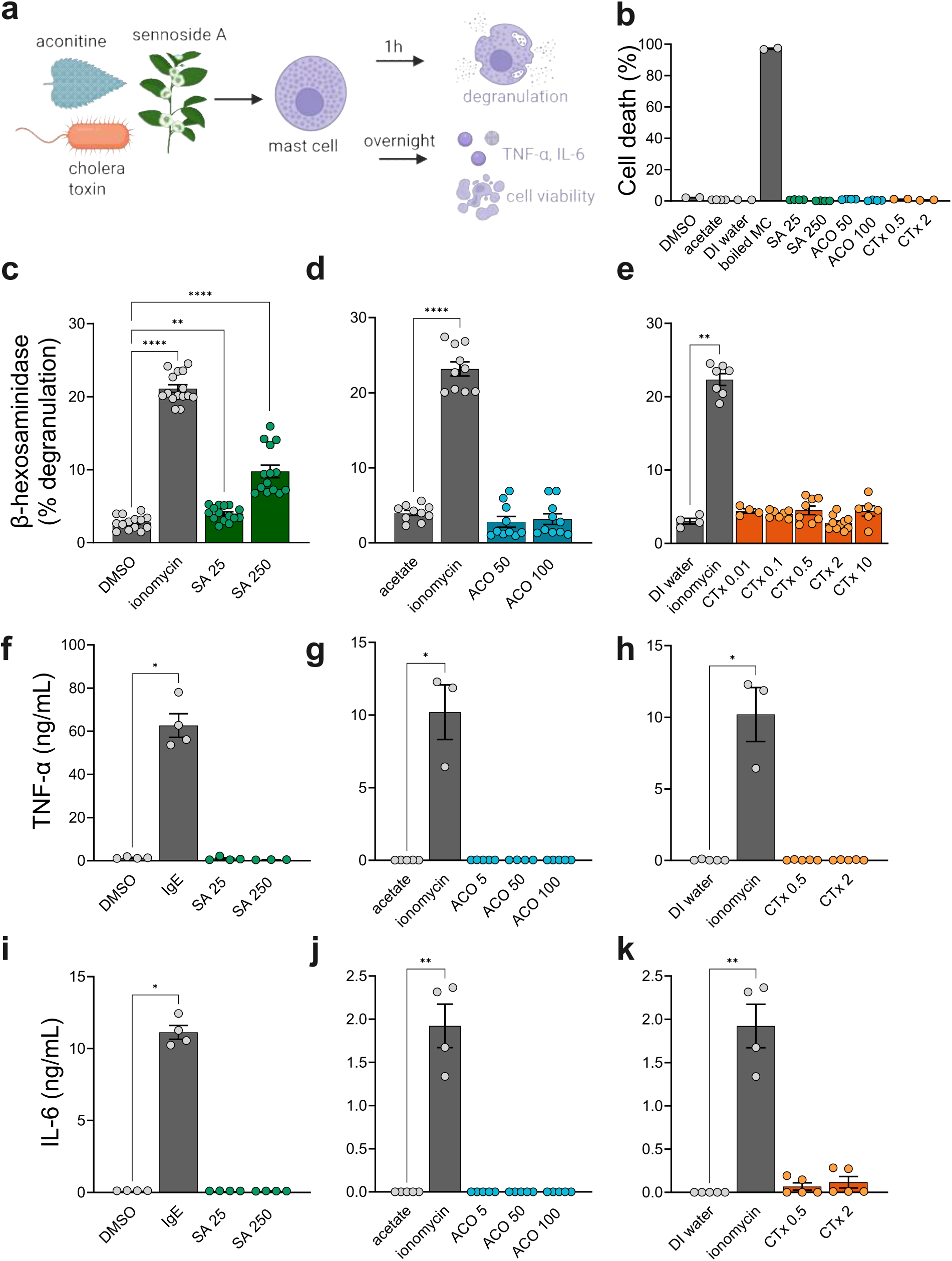
Sennoside A selectively induces mast cell degranulation without reducing mast cell viability. **a,** Experimental schematic. Mast cells were exposed to sennoside A, aconitine, or cholera toxin for 1 h prior to quantification of β-hexosaminidase release, and for 24 h before measuring cell viability and the secretion of TNF-α and IL-6. **b,** Cell viability as a percentage of Zombie Yellow⁺ cells by flow cytometry. **c–e,** Percent β-hexosaminidase release from mast cells acutely exposed to varying doses of toxins. Ionomycin and vehicle controls (1% DMSO, 0.5 M acetate, DI water) were used as positive and negative controls, respectively. **f–k**, Supernatant concentrations of TNF-α and IL-6 quantified by ELISA. **b–k,** Graphs show mean ± s.e.m. *p < 0.05, **p < 0.01, ****p < 0.0001. Mann–Whitney U. Each panel is representative of at least two independent experiments. **a,** created with <u>Biorender.com</u>.

We next measured β-hexosaminidase release as a readout of mast cell degranulation after 1 h of exposure (Fig. 1a). Sennoside A increased β-hexosaminidase release relative to vehicle, with a larger response at 250 μg/mL (Fig. 1c). The response remained lower than that elicited by ionomycin: ionomycin induced 5.3-fold and 2.2-fold more β-hexosaminidase release than 25 and 250 μg/mL sennoside A, respectively. In contrast, neither aconitine nor cholera toxin significantly increased degranulation at the concentrations tested (Fig. 1d, e).

We also measured TNF-α and IL-6 secretion after toxin exposure (Fig. 1a). Neither cytokine was significantly increased by sennoside A, aconitine, or cholera toxin compared with the corresponding vehicle controls (Fig. 1f–k). Thus, under these conditions, sennoside A elicited degranulation without a detectable increase in these two inflammatory cytokines.

These results indicate that mast cell responses were compound-selective rather than a nonspecific consequence of chemical exposure. Because sennoside A induced degranulation without measurable loss of viability, we next tested whether it altered IgE-dependent mast cell activation.

### Sennoside A suppresses IgE-mediated mast cell activation in vitro

To test the effect of these compounds on allergic activation, BMMCs were exposed overnight to sennoside A, aconitine, or cholera toxin during sensitization with anti-DNP IgE and were then challenged with DNP-BSA (Fig. 2a). Sennoside A markedly reduced antigen-induced degranulation at both concentrations tested. β-hexosaminidase release was 7.3-fold lower with 25 μg/mL sennoside A and 5.7-fold lower with 250 μg/mL sennoside A than in the IgE/DNP-BSA positive-control condition (Fig. 2b). Aconitine and cholera toxin did not significantly alter antigen-induced degranulation (Fig. 2c, d).

**Figure 2.**
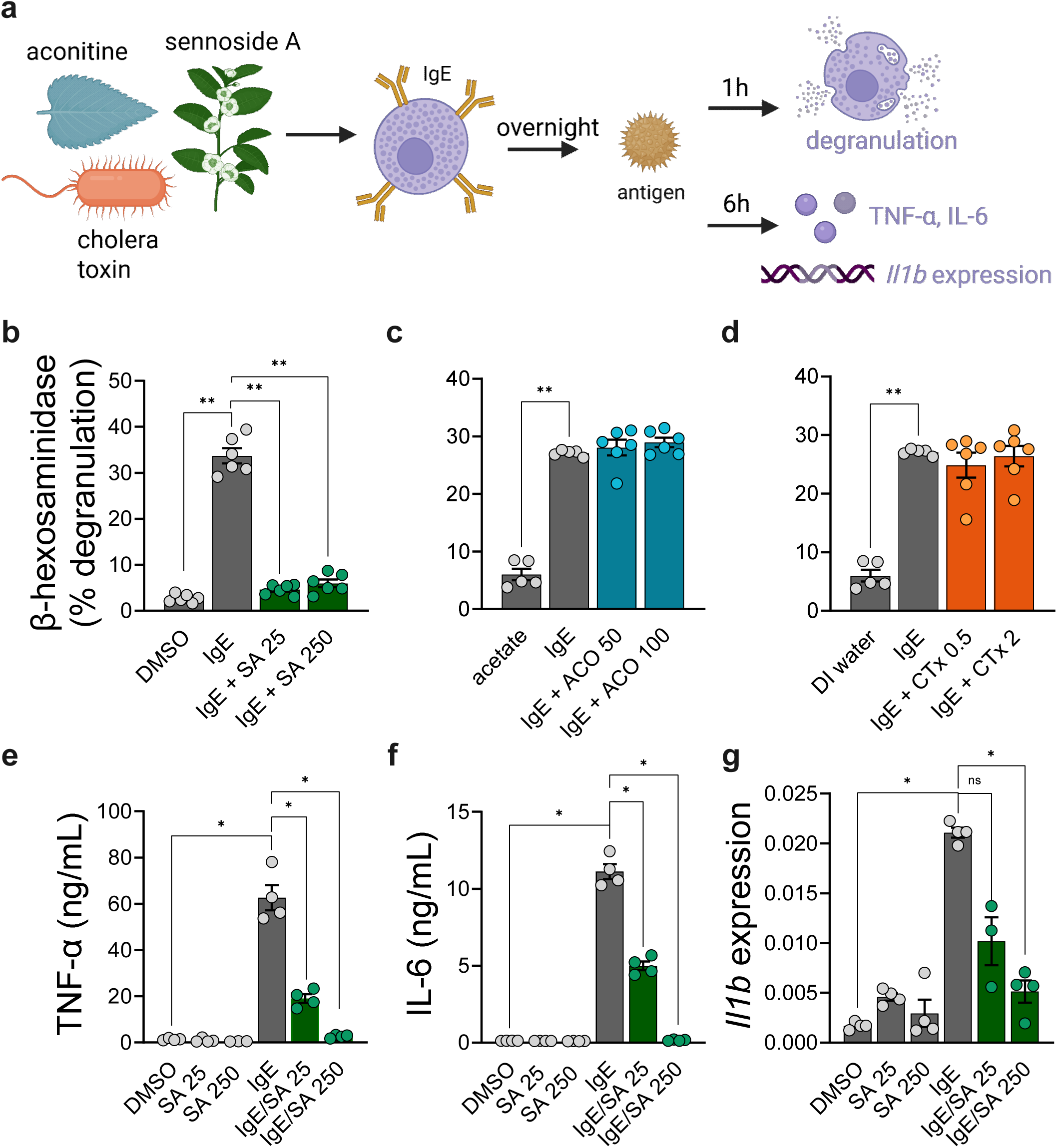
Sennoside A suppresses IgE-mediated mast cell activation *in vitro*. **a,** Experimental schematic. Mast cells were pretreated overnight with sennoside A, aconitine, or cholera toxin in the presence of anti-DNP IgE prior to challenge with DNP-BSA antigen. β-hexosaminidase release was quantified 1 h following antigen challenge, while TNF-α and IL-6 secretion, as well as IL-1β expression, were assessed 6 h following antigen challenge. **b–d,** Percent β-hexosaminidase release from IgE-sensitized mast cells following acute DNP-BSA stimulation. Anti-DNP IgE/DNP-BSA and vehicle controls (1% DMSO, 0.5 M acetate, DI water) were used as positive and negative controls, respectively. **e–f, Supernatant** concentrations of TNF-α and IL-6 quantified by ELISA. **g,** Expression of *Il1b* relative to *Rpl13a* gene expression quantified by qPCR. **b–g,** Graphs show mean±s.e.m. *p<0.05, **p<0.01. Mann–Whitney U. Each panel is representative of at least two independent experiments. **a**, created with <u>Biorender.com</u>.

Sennoside A also reduced TNF-α and IL-6 secretion after IgE-dependent activation (Fig. 2e, f). In addition, 250 μg/mL sennoside A significantly reduced *Il1b* expression after antigen challenge (Fig. 2g). These effects were not observed with aconitine or cholera toxin under the conditions tested.

Thus, sennoside A had distinct effects depending on the activation context: it promoted modest degranulation in resting BMMCs but strongly suppressed degranulation and inflammatory mediator production during IgE-dependent activation. We next examined the physiological effects of oral sennoside A and its impact in a mouse model of food allergy.

### Acute oral sennoside A induces hypothermia, diarrhea, and colonic inflammatory gene expression

To characterize the acute response to oral sennoside A, male and female BALB/c mice received a single intragastric dose of 10 or 30 mg/kg during the active phase (ZT14) and were monitored for 6 h (Fig. 3a). Mice given 30 mg/kg, but not 10 mg/kg, developed hypothermia and diarrhea (Fig. 3b–d).

**Figure 3.**
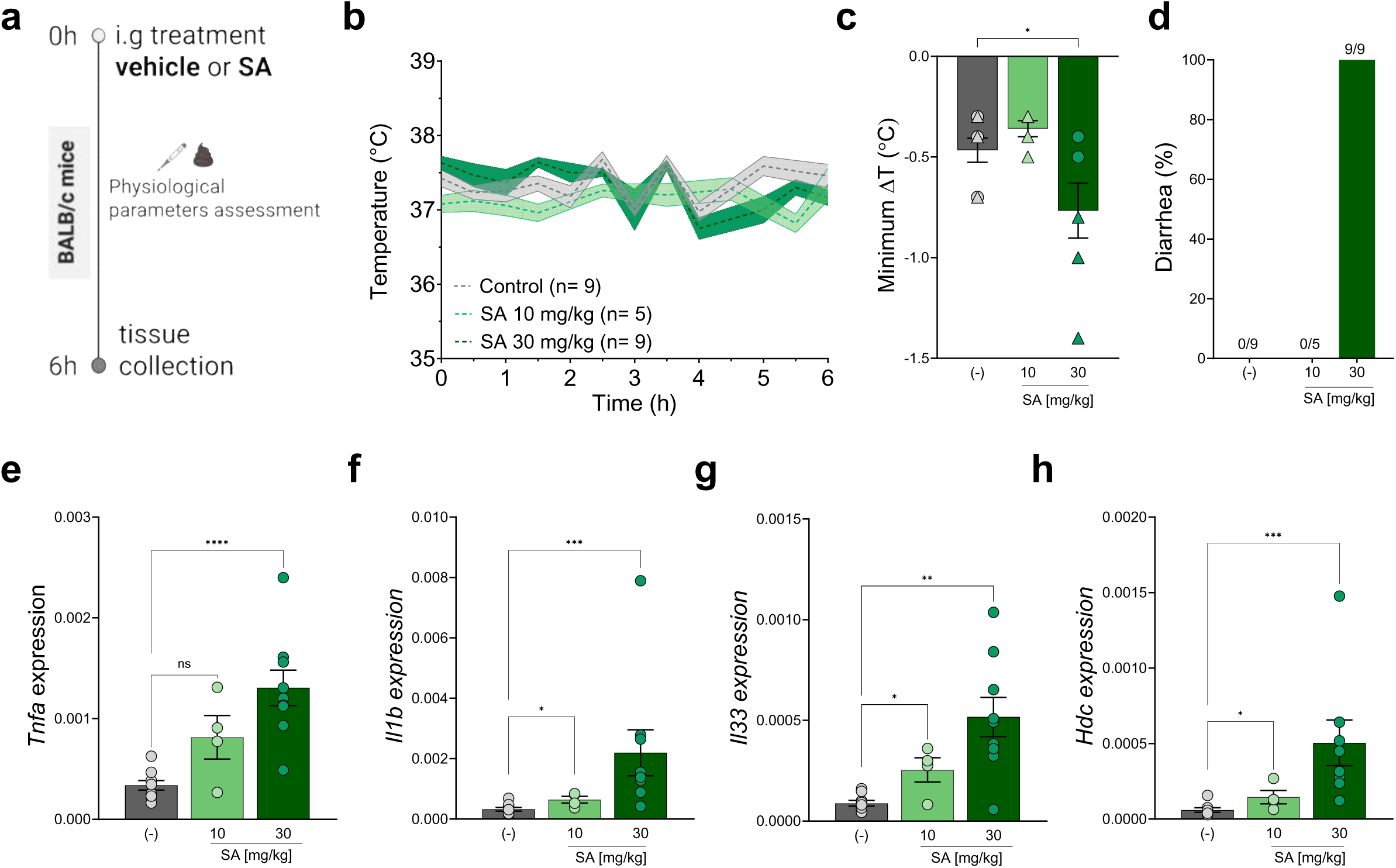
Acute oral sennoside A induces hypothermia, diarrhea, and colonic inflammatory gene expression. **a,** Schematic of the experimental protocol for acute sennoside A exposure and assessment of physiological responses. **b,** Rectal temperature over time following oral sennoside A exposure. **c,** Maximum decrease in rectal temperature (n= 5-9 mice per group). **d,** Diarrhea occurrence (n= 5-9 mice per group). **e-h,** Colonic expression of **e,** tumor necrosis factor alpha (*Tnfa*), **f,** interleukin-1 beta (*Il1b*), **g,** interleukin-33 (*Il33*), **h,** histidine decarboxylase (*Hdc*) relative to *Rpl13a* gene expression quantified by qPCR. **e-h,** Graphs show mean±s.e.m. *p<0.05, **p<0.01, ***p < 0.001, ****p < 0.0001. Mann–Whitney U. Each panel is representative of at least two independent experiments. **a**, created with <u>Biorender.com</u>.

Because the laxative activity of sennoside A depends on metabolism by the colonic microbiota (23), we assessed gene expression in the large intestine 6 h after administration. Sennoside A treatment was associated with increased expression of *Tnfa*, *Il1b*, *Il33*, and *Hdc* (Fig. 3e–h). These data show that acute oral sennoside A can produce systemic physiological changes together with a rapid intestinal transcriptional response.

### Sennoside A attenuates allergen-induced hypothermia and reduces circulating IgE and OVA-specific IgG1

We next asked whether the inhibitory effect of sennoside A on IgE-dependent mast cell activation *in vitro* was accompanied by reduced allergic responses *in vivo*. Female BALB/c mice were sensitized with OVA and alum on days 0 and 7 and then challenged orally with OVA five times (Fig. 4a) (6). Beginning after sensitization, mice received sennoside A (20 mg/kg, i.g.) 24 h before each oral OVA challenge. This intermediate dose was selected to fall between 10 mg/kg, which produced little acute physiological response, and 30 mg/kg, which induced hypothermia and diarrhea. Tissues and serum were collected after the fifth challenge.

**Figure 4.**
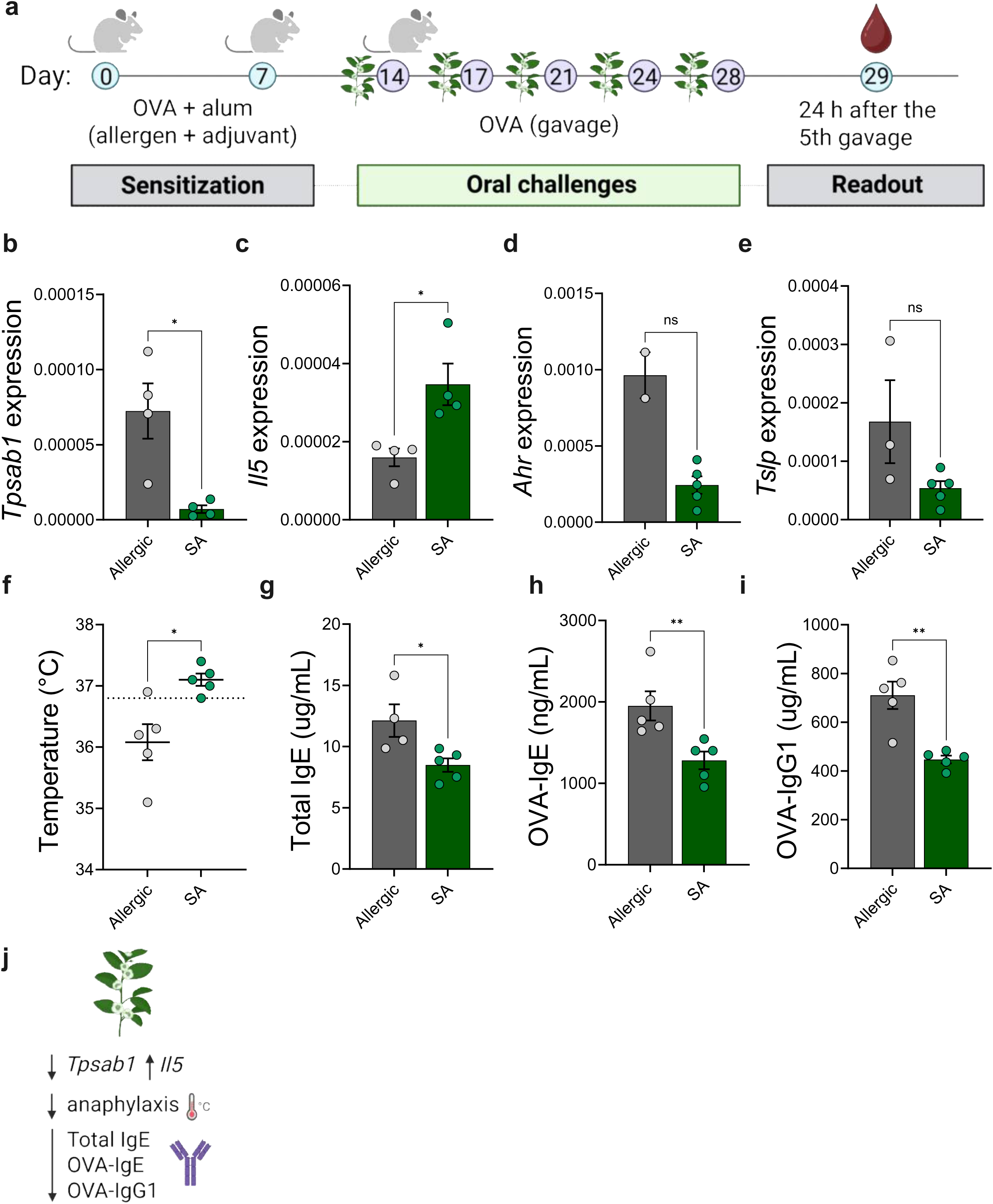
Sennoside A attenuates allergen-induced hypothermia and reduces circulating IgE and OVA-specific IgG1. **a,** Experimental schematic. BALB/c mice were sensitized by subcutaneous injections of ovalbumin (OVA) and the adjuvant aluminum hydroxide (alum) on days 0 and 7. After sensitization, mice received five intragastric (i.g.) challenges with 40 mg of OVA. Sennoside A (20 mg/kg) was administered i.g. 24 h before each OVA challenge. Rectal temperature was measured immediately following the fifth OVA gavage, and tissues and blood were collected 24 h later. **b–e,** Duodenal expression of **b**, tryptase alpha/beta 1 (*Tpsab1),* **c,** interleukin-5 (*Il5),* **d,** aryl hydrocarbon receptor (*Ahr*), and **e,** thymic stromal lymphopoietin (*Tslp*) relative to *Rpl13a* gene expression quantified by qPCR. **f,** Change in rectal temperature (℃) measured 1 h following the fifth OVA challenge. The dotted line represents the average rectal temperature of both groups at 0 h. **g–h,** Serum total IgE, serum OVA-specific IgE, and serum OVA-specific IgG1 concentrations quantified by ELISA. **j,** Summary of findings. **b–i,** Graphs show mean±s.e.m. *p<0.05, **p<0.01. Mann–Whitney U. Each panel is representative of at least two independent experiments. **a, i,** created with <u>Biorender.com</u>.

In the duodenum, sennoside A-treated allergic mice had lower expression of *Tpsab1*, a mast cell-enriched tryptase gene, than untreated allergic controls (Fig. 4b). *Il5* expression was increased (Fig. 4c), whereas *Ahr* and *Tslp* were not significantly different between groups (Fig. 4d,e).

We next assessed allergen-induced hypothermia, a feature of systemic reactions in this model. One hour after the fifth OVA challenge, untreated allergic mice showed an average decrease in rectal temperature of approximately 0.75°C, whereas sennoside A-treated mice maintained temperatures closer to baseline (Fig. 4f).

Sennoside A treatment was also associated with lower circulating allergic antibodies. Total IgE was reduced by approximately 1.4-fold, OVA-specific IgE by approximately 1.6-fold, and OVA-specific IgG1 by approximately 1.6-fold relative to untreated allergic controls (Fig. 4g–i).

Together, these results show that sennoside A administration during the allergen-challenge phase attenuated allergen-induced hypothermia and was associated with lower circulating IgE and OVA-specific IgG1, lower duodenal *Tpsab1* expression, and increased *Il5* expression. Because sennoside A treatment began after sensitization, these findings indicate effects on responses occurring during repeated allergen challenge rather than on the initial sensitization step.

## DISCUSSION

The main finding of this study is that sennoside A exerts distinct effects on mast cells depending on their activation state. In resting bone marrow-derived mast cells, sennoside A induced modest degranulation without detectable TNF-α or IL-6 secretion. In contrast, during IgE-dependent activation, sennoside A markedly suppressed antigen-induced degranulation, TNF-α and IL-6 secretion, and expression of IL-1β. This inhibitory effect was accompanied by attenuation of several features of experimental food allergy *in vivo*, including allergen-induced hypothermia and lower circulating total and antigen-specific IgE. These findings identify sennoside A as a previously unrecognized modulator of mast cell function and show that the effect of a naturally occurring bioactive compound can depend strongly on the underlying immune context.

The suppression of IgE-mediated mast cell activation by sennoside A is particularly relevant to mast cell biology in allergic disease. Crosslinking of FcεRI initiates a coordinated signaling cascade that drives rapid granule exocytosis followed by production of cytokines, lipid mediators, and other inflammatory molecules (24). Sennoside A reduced both β-hexosaminidase release and production of TNF-α and IL-6, indicating that its inhibitory effect is not restricted to a single mast cell output. The molecular target responsible for this effect remains unknown. Potential sites of regulation include proximal FcεRI signaling, PLCγ-dependent calcium mobilization, MAPK signaling, and transcriptional pathways such as NF-κB. Plant-derived compounds have previously been shown to interfere with mast cell activation at several of these levels. For example, quercetin and kaempferol suppress IgE-mediated β-hexosaminidase release and cytokine production together with reduced phosphorylation of Syk, PLCγ, ERK, p38, and JNK (25). Tetrahydrocurcumin similarly inhibits IgE-mediated mast cell degranulation and cytokine production and can also suppress ionomycin- and MRGPRX2-mediated activation (26). These studies illustrate that naturally occurring compounds can regulate mast cell responses through multiple signaling nodes and provide candidate pathways through which sennoside A may act.

Studies in other immune cell populations provide additional clues. In macrophages, sennoside A has been reported to suppress NF-κB-dependent NLRP3 priming and inflammasome activation and to directly inhibit caspase-1 activity (27). NF-κB regulation could contribute to the reduced TNF-α, IL-6, and IL-1β observed in our experiments. NLRP3 may also be relevant to mast cell degranulation independently of its canonical role in inflammasome activation. Mencarelli and colleagues reported that NLRP3 facilitates granule trafficking and exocytosis during IgE-mediated mast cell activation (28). Whether either pathway contributes to the inhibitory effect of sennoside A remains to be determined. Direct examination of FcεRI signaling, calcium mobilization, MAPK and NF-κB activation, and NLRP3 function will be necessary to define the molecular basis of this response.

Interestingly, the inhibitory effect of sennoside A during IgE-dependent activation contrasted with its ability to induce degranulation in otherwise unstimulated mast cells. Sennoside A-induced degranulation occurred without a corresponding increase in TNF-α or IL-6, suggesting that the response is qualitatively different from canonical FcεRI-mediated activation. Dissociation between degranulation and cytokine production has also been observed in response to other mast cell stimuli. The *Staphylococcus aureus* δ-toxin, for example, directly induces mast cell degranulation through mechanisms involving PI3K signaling and calcium influx without compromising cell viability (19), whereas the hemolytic pigment produced by Group B *Streptococcus* induces degranulation together with lipid mediator and inflammatory cytokine production and cellular injury (22). Plant-derived compounds can likewise produce distinct mast cell phenotypes. The lectin ArtinM induces IL-6 production without degranulation at lower concentrations but stimulates both responses at higher concentrations (29). These observations, together with our findings, indicate that mast cells do not mount a uniform response to chemically diverse environmental compounds. Instead, individual compounds can selectively engage different components of the mast cell activation program.

The *in vivo* food allergy experiments are consistent with an inhibitory effect of sennoside A on allergic inflammation, although they do not establish mast cells as its only cellular target. Sennoside A treatment attenuated the allergen-induced decrease in body temperature, a hallmark of the systemic response to oral allergen challenge that depends on IgE-FcεRI signaling and intestinal mast cell activation (17). Treatment was also associated with reduced duodenal *Tpsab1* expression. Because *Tpsab1* encodes mast cell tryptase, this finding is consistent with an alteration in intestinal mast cell abundance, phenotype, or activation state, but tissue transcript abundance alone cannot distinguish among these possibilities. The *in vitro* experiments provide complementary evidence that sennoside A can directly suppress IgE-dependent mast cell responses, supporting the possibility that mast cell inhibition contributes to the *in vivo* phenotype.

Sennoside A also reduced circulating total IgE, OVA-specific IgE, and OVA-specific IgG1. These changes suggest that its effects *in vivo* are not necessarily restricted to mast cells and may extend to pathways regulating the humoral immune response. Because sennoside A treatment was initiated after systemic sensitization and administered before oral allergen challenges, our experiments do not establish that sennoside A prevents initial allergic sensitization. Rather, they indicate that repeated treatment during the challenge phase is associated with lower antibody concentrations at the end of the experimental protocol. Determining whether this reflects altered B cell responses, T cell help, intestinal antigen handling, or secondary effects of reduced inflammation will require further study.

A second unexpected feature of the allergic model was the increase in intestinal Il5 expression following sennoside A treatment. IL-5 is best known for promoting eosinophil development, recruitment, and survival and is commonly associated with type 2 inflammation (30,31). However, eosinophils can also contribute to tissue protection and immune regulation in the gastrointestinal tract (32–34). Recent studies have shown that eosinophils can reduce allergic diarrhea and mortality from anaphylactic shock in experimental food allergy (35). IL-5 has also been implicated in supporting mucosal IgA responses and intestinal barrier function (36). The increased *Il5* expression observed here could therefore reflect enhanced eosinophilic inflammation, a compensatory tissue response, or a protective mucosal program. Measurements of intestinal eosinophil abundance and activation, together with assessment of mucosal antibody responses, will be necessary to determine the significance of this finding.

The protective phenotype observed in the food allergy model is consistent with anti-inflammatory effects of sennoside A reported in other experimental settings. Oral sennoside A has been shown to reduce circulating and tissue inflammatory mediators in models of metabolic inflammation (14). Other plant-derived compounds can also suppress experimental allergic disease. Tetrahydrocurcumin attenuates the anaphylactic decrease in body temperature in a mouse model of peanut allergy, while curcumin reduces allergic diarrhea in an OVA-induced model (26,37). Notably, these compounds may achieve similar physiological outcomes through different effects on mast cells. Tetrahydrocurcumin inhibits mast cell activation without reducing cell viability, whereas curcumin has been reported to induce mast cell apoptosis (26,37). In our experiments, sennoside A suppressed allergic mast cell activation without detectable cytotoxicity, supporting regulation of mast cell function rather than loss of mast cells as the explanation for its effects *in vitro*.

The absence of cytotoxicity was also notable in the initial comparison of sennoside A, aconitine, and cholera toxin. None of the three compounds reduced mast cell viability at the concentrations and exposure times tested. This should not be interpreted as evidence that mast cells are broadly resistant to toxic compounds. Several microbial and nonmicrobial toxicants, including *Clostridioides difficile* toxin A, *Pseudomonas aeruginosa* exotoxin A, selected metal ions, and the plant-derived monoterpene thymol, have been reported to induce mast cell injury or death (38–41). Thus, the lack of cytotoxicity observed here most likely reflects compound-specific differences in mast cell susceptibility. Likewise, neither aconitine nor cholera toxin induced detectable degranulation or TNF-α or IL-6 production under our experimental conditions. These negative responses reinforce the idea that mast cell activation is selective rather than a general consequence of exposure to biologically active or toxic compounds.

The acute effects of orally administered sennoside A further illustrate the context-dependent biology of this compound. Sennoside A is metabolized by the colonic microbiota to rheinanthrone, which promotes intestinal secretion and motility through mechanisms that include prostaglandin E2 production and altered epithelial water transport (23,42). This pathway underlies the established laxative activity of sennosides and is consistent with the diarrhea observed in our study. In contrast, the hypothermia and inflammatory transcriptional response were less expected. To our knowledge, acute hypothermia following oral administration of sennoside A has not been previously described. Hypothermia can occur as part of the physiological response to toxic insults in laboratory mammals and has been proposed to reduce metabolic demand and limit injury under some conditions (43), although the mechanism responsible for the response to sennoside A remains unknown.

Acute ingestion of sennoside A also increased colonic expression of TNF-α, IL-1β, IL-33, and histidine decarboxylase. Increased *Tnfa* and *Il1b* is consistent with activation of local inflammatory pathways, whereas *Il33*, which encodes an epithelial alarmin associated with type 2 immunity, may reflect epithelial stress or mucosal immune activation (44–46). Increased *Hdc* suggests enhanced local capacity for histamine synthesis and is particularly interesting given the ability of sennoside A to induce mast cell degranulation *in vitro*. Histamine is a major mediator of allergic responses and can alter vascular tone, vascular permeability, and smooth muscle function through its actions on multiple target tissues (47). More broadly, it has been proposed that immune-mediated changes in blood flow during responses to harmful foods could contribute to postabsorptive toxin defense by altering the distribution of circulating toxins and thereby limiting exposure of vulnerable tissues (7). Although our experiments do not establish increased histamine production or a protective function for this response, the induction of *Hdc* raises the possibility that histaminergic signaling forms part of the physiological response to ingested sennoside A. The cellular source and functional consequence of increased *Hdc* expression therefore remain to be determined. Moreover, because orally administered sennoside A is metabolized by the intestinal microbiota to rheinanthrone, the *in vivo* response could be mediated by intact sennoside A, its metabolites, or both. The direct response of cultured mast cells to sennoside A indicates that microbial conversion is not required for all its biological effects; however, direct comparison of sennoside A and rheinanthrone will be necessary to distinguish their respective contributions *in vivo*.

Several limitations should be considered when interpreting these findings. Our analysis of mast cell activation focused on β-hexosaminidase release and TNF-α and IL-6 production. Although these are established measures of degranulation and inflammatory activation, they capture only a subset of mast cell responses. Broader transcriptomic, proteomic, or lipid-mediator profiling could reveal additional pathways regulated by sennoside A. We also examined only IgE/FcεRI-mediated allergic activation; whether sennoside A similarly affects IgE-independent mast cell activation remains unknown. In addition, the *in vitro* experiments were performed using murine bone marrow-derived mast cells. Because mast cell phenotypes differ across tissues and species, studies using intestinal mast cells and primary human mast cells will be important for determining the broader relevance of these observations. Finally, although the food allergy model used here is dependent on IgE and mast cell effector responses (6,16,17), our experiments do not directly establish that mast cells mediate the protective effects of sennoside A *in vivo*. Reduced *Tpsab1* expression and attenuation of allergen-induced hypothermia are consistent with altered mast cell responses, and the *in vitro* experiments demonstrate that sennoside A can act directly on mast cells. Nevertheless, changes in circulating IgE and IgG1, intestinal *Il5*, and other immune pathways indicate that additional cell populations are likely to contribute. Establishing the requirement for mast cells *in vivo* and identifying the molecular target of sennoside A will be important next steps.

Mast cells can couple immune recognition of ingested antigens to organism-level responses, including anaphylaxis and antigen avoidance (6,17). Here, we extend this concept by showing that a plant-derived dietary compound can itself alter mast cell function, with markedly different consequences depending on the underlying activation state. Sennoside A promoted modest degranulation in resting mast cells yet suppressed IgE-dependent degranulation and inflammatory mediator production during allergic activation. *In vivo*, repeated oral treatment attenuated several features of experimental food allergy, whereas acute exposure elicited distinct physiological and intestinal inflammatory responses. These observations expand the range of naturally occurring compounds known to modulate mast cell biology and highlight the importance of immune context in determining how mast cells respond to environmental and dietary bioactive molecules.

## METHODS

### Mice

BALB/c mice (6-10 weeks old; Charles River Laboratories, USA) were used for all experiments. All animals were housed in temperature- and humidity-controlled rooms, in a 12/12h light/dark cycle, with lights off from 7:00 AM to 7:00 PM (ZT1-12). Food and water were provided ad libitum. Experiments were conducted in the animal facility at the Biodesign Institute, Arizona State University (Arizona, USA). All animal care and experimentation were approved by the Institutional Animal Care and Use Committee of Arizona State University and consistent with the National Institutes of Health, USA, guidelines. All procedures were approved by the Institutional Animal Care and Use Committee at Arizona State University (protocols #21-1864R, #24-2033R, #26-2184)

### Key Resources Table

| REAGENT or RESOURCE | SOURCE | CATALOG No. |
| --- | --- | --- |
| **Antibodies** |  |  |
| anti-CD16/CD32 (Fc block) (dilution 1:200) | BD Pharmingen | 553142 |
| OVA-specific IgE capture antibody | BD Pharmingen | 553413 |
| Total IgE capture antibody | BD Pharmingen | 553413 |
| Total IgE detection antibody | BD Pharmingen | 553414 |
| OVA-specific IgG1 detection antibody | BD Pharmingen | 553441 |
| FceR1 (clone: MAR-1) (dilution 1:200) | Invitrogen | 12-5898-82 |
| CD117 (Ckit) (clone: 2B8) (dilution 1:200) | Invitrogen | 48-1171-82 |
| **Standards for ELISAs** |  |  |
| OVA-specific IgE standard (anti-OVA IgE) | Bio-Rad | MCA2259 |
| Total IgE standard | BD Pharmingen | 557080 |
| IgG1 standard | BD Pharmingen | 557273 |
| **Chemicals** |  |  |
| Cholera toxin | Millipore Sigma | 227036 |
| Aconitine | Millipore Sigma | 44787 |
| Sennoside A | Millipore Sigma | 68909 |
| Ionomycin | Millipore Sigma | 407951 |
| Anti-Dinitrophenyl (DNP) IgE | Millipore Sigma | D8406 |
| DNP-BSA | Millipore Sigma | 324101 |
| Triton X-100 | Sigma-Aldrich | Cat# X100-100ML |
| Dulbecco's Phosphate Buffered Saline (1X) | Corning | Cat# 21-040-CV |
| Glycine | Sigma-Aldrich | Cat# 50046-50G |
| RNA Stat | Amsbio | Cat# CS-502-500 |
| Chloroform | Sigma-Aldrich | Cat#650498 |
| Oligo(dT) primer | Millipore Sigma | Cat#69896 |
| PerfeCTa® SYBR® Green SuperMix, Low ROX™ | QuantaBio | Cat#95056-02K |
| Phosphate-buffered saline (PBS, 1X), sterile-filtered | ThermoFisher Scientific | Cat# J61196.AP |
| Fetal Bovine Serum | PeakSerum | Cat# PS-FB3 |
| Trypan Blue | Thermo Fisher | Cat# 15250061 |
| grade III OVA | Sigma-Aldrich | A5378 |
| Alhydrogel 2% (aluminium Hydroxide Gel) | InvivoGen | Vac-alu-250 |
| Endo Grade® Ovalbumin, 25 mg | BioVendor | LET0027 |
| **Commercial assays** |  |  |
| TMB Substrate Reagent Set | BD Biosciences | 555214 |
| BD Streptavidin HRP Solution | BD | 554066 |
| RNeasy Mini Kit (250) | Qiagen | Cat#74106 |
| SMART® MMLV Reverse Transcriptase | Takara | Cat#639524 |
| **Experimental model: Organisms/strains** |  |  |
| Mouse: BALB/c<br>In-house bred | Charles River Laboratories | Strain code: 028 |
| **Software** |  |  |
| GraphPad Prism version 11.1 | GraphPad Software |  |
| Adobe Illustrator version 27.3.1 | Adobe Inc. |  |
| FlowJo version 10.9.0 | Becton, Dickinson and Company |  |
| **Equipment** |  |  |
| Attune NxT Flow Cytometer | ThermoFisher Scientific |  |
| Revolve fluorescent microscope, model 2-K1214 | ECHO, San Diego, CA |  |
| ViiA 7 Real-Time PCR System | ThermoFisher Scientific |  |
| SimpliAmp™ Thermal Cycle | Applied Biosystems, Thermo Fisher Scientific |  |

### Bone marrow–derived mast cells

Bone marrow–derived mast cells (BMMCs) were generated by culturing bone marrow cells isolated from mice. Mice were euthanized and bone marrow femurs and tibias were isolated under sterile conditions. Both ends of each bone were cut, and the bones were flushed with high-glucose DMEM (Sigma-Aldrich) supplemented with 10% FBS (Peak Serum), 25 mM HEPES (VWR), 1% penicillin/streptomycin (VWR), 1% L-glutamine (SAFC), 1% sodium pyruvate (Corning), 1% MEM non-essential amino acids (Corning), and 50 μM β-mercaptoethanol (Sigma-Aldrich) using a 10 mL syringe. Marrow was transferred to a 50 mL conical tube after straining with a 70 µm cell strainer. Cells were centrifuged (250 g, 5 min), supernatant was discarded, and the pellet was resuspended in complete media further supplemented with 20 ng/mL recombinant murine IL-3 and 25 ng/mL recombinant murine stem cell factor (SCF) (PeproTech). Cells were cultured for 6 weeks, with media replaced twice weekly.

### Cell culture and treatments

For culture changes, cells were centrifuged (250 g, 5 min), supernatant was aspirated, and cells were resuspended in complete media and counted. All stimulations, pre-treatments are described in the main text and figure legends.

### Flow cytometry

#### Mast cell maturity and purity

BMMC purity and maturity were assessed by flow cytometry using anti-mouse CD117 (c-Kit) (Invitrogen, 48-1171-82, clone: 2B8, 1:200), and FcεRI antibodies (Invitrogen, 12-5898-82, clone: MAR-1, 1:200). Cells (3 × 10⁵) were washed in FACS buffer (2% FBS in PBS) and stained with 1 μg/mL of each antibody for 30 min at 4°C. After washing, cells were fixed with 1% paraformaldehyde and analyzed on Attune NxT Flow Cytometer (ThermoFisher Scientific, B2R3V6Y3). Data were analyzed using FlowJo™ Software (Becton, Dickinson, and Company). Cells were gated to identify single cells, then single-color controls using stained cells were established for positive detection of CD117 and FcεRI. Cultures with >90% CD117⁺ FcεRI⁺ cells were used for experiments.

#### Viability assay

Cell viability was assessed using Zombie Yellow viability dye (BioLegend 423104) according to the manufacturer’s protocol. Cells (1 × 10⁵) were incubated with the dye for 20 min at room temperature (RT), washed in FACS buffer, and analyzed by flow cytometry. The gating strategy was drawn from negative controls, first gating to isolate cells from debris, followed by single cells from doublets, and then viable cells (Zombie Yellow-negative).

### Mast cell degranulation assay

β-hexosaminidase degranulation assay was performed as previously described (48). Briefly, BMMCs were plated overnight at 5 × 10⁵ cells per well in 24-well plates in complete medium. Cells were washed three times with Tyrode’s buffer and resuspended in 40 μL Tyrode’s buffer with indicated stimuli. Following stimulation for 1 h at 37°C, cells were centrifuged (450 g, 5 min, 4°C), and 20 μL of supernatant was collected. Cell pellets were lysed with 0.5% Triton X-100 in PBS, centrifuged, and 20 μL lysate was collected. Supernatants and lysates were incubated with 50 μL p-nitrophenyl-N-acetyl-β-D-glucosamine (1.3 mg/mL in 0.1 M sodium citrate, pH 4.5) for 1 h at 37°C. Reactions were stopped with 150 μL 0.4 M glycine buffer (pH 10.7), and absorbance was measured at 405 nm. Degranulation (%) was calculated as: degranulation (%) = (supernatant / [supernatant + lysate]) × 100

### Toxin exposure

Cells were stimulated with cholera toxin (0.01–10 μg/mL), sennoside A (25 and 250 μg/mL), or aconitine (50 and 100 μg/mL,) for 1 h. Briefly, cholera toxin was reconstituted in deionized water and stored at 4°C. Aconitine was reconstituted in 0.1 M acetate buffer and stored at -20°C. Sennoside A was reconstituted in DMSO and stored at -20°C. Therefore, vehicle controls included deionized water (cholera toxin), DMSO (sennoside A), and acetate buffer (aconitine). Ionomycin (1 μM) was used as a positive control. Degranulation was quantified as described above.

For IgE-dependent activation, cells were incubated overnight with toxins in the presence or absence of 2 μg/mL anti-DNP IgE. On the following day, cells were stimulated with 25 μg/mL DNP-BSA prior to degranulation analysis.

To quantify cytokine secretion, BMMCs (4 × 10⁶) were plated overnight in 12-well plates in a final volume of 1.5 mL. Cells were treated with toxins with or without IgE sensitization and stimulated with either ionomycin (1 μM) or DNP-BSA (25 μg/mL) for 6 h.

### Cytokine quantification

Supernatant levels of TNF-α and IL-6 were determined by sandwich enzyme-linked immunosorbent assay (ELISA). For TNF-α, ELISA-grade plates (Thermo Fisher, 442404) were coated with 2 μg/mL of anti-mouse TNF-α (eBioscience, 13-7423-85) in 0.1M sodium carbonate buffer (pH 9.5) overnight at 4°C. Plates were blocked with 1% bovine serum albumin (Sigma-Aldrich, A7030) at RT for 1 h. Culture supernatant (0.1 mL) was then incubated at RT for 2 h. Purified mouse TNF-α (Invitrogen, 29-8321-65) was used as a standard curve, with the highest concentration being 4 ng/mL followed by two-fold dilutions to 0.0002 ng/mL. After sample incubation, biotin-conjugated TNF-α detection antibodies (eBioscience, 13-7341-85) at 1 μg/mL were incubated at RT for 1 h. Next, 1:1000-fold dilution of HRP-conjugated streptavidin (BD Biosciences, 554066) was incubated at RT for 30 min. Plates were then incubated with TMB substrate reagent (BD Biosciences, 555214) in the dark at RT, with the color checked every 2 min. Reactions were stopped with 2.5M sulfuric acid (Ricca, UN2796) at the appearance of a two-fold dilution on the standard curve and absorbance at 450nm was read immediately using a plate reader (Varioskan Lux, Thermo Scientific, Waltham, MA). Between each step, plates were washed 3-7 times with 0.05% Tween-20 (Sigma-Aldrich, P1379) in PBS. Supernatant levels of IL-6 were measured following the same method. Plates were coated with 2 μg/mL of anti-mouse IL-6 (eBioscience, 14-7061-85) used as the capture antibody, and 1 μg/mL of biotin-conjugated IL-6 antibodies (BD Biosciences, 554402) used for detection. Purified mouse IL-6 (R&D Systems, DY406-05) was used as a standard curve, with the highest concentration being 16 ng/mL followed by two-fold dilutions to 0.0078 ng/mL.

### RNA isolation and qPCR

Following stimulation, cells were collected in RNA-STAT-60 (Gentuar, CS-502-500) and immediately stored at -70°C until processed for RNA extraction. RNA was extracted using the RNeasy Mini-Kit (Qiagen, 74004) according to the manufacturer’s specifications.

Briefly, cells were pelleted at 250 g for 5 minutes, and supernatant was removed. Cells were washed with 1 mL cold 1x PBS was added and pelleted again. PBS was discarded and 0.7 mL of RNA-STAT-60 was added for 1 minute and vortexed for 45 seconds. Chloroform (0.14 mL) was added and samples were left on ice for 5 minutes. Samples were centrifuged at maximum rcf for 10 minutes at 4 °C. The top aqueous layer (RNA) was collected (maximum of 0.35 mL) and transferred to an eppendorf tube. Cold 70% ethanol (0.35 mL) was added to the eppendorf tube and samples were left on ice for 5 minutes before transferring to a spin column with a 2 ml collection tube. Samples were centrifuged at 11,000 g for 30 seconds and flow-through was discarded.

DNase digestion was performed by adding 0.35 mL of RW1 buffer to the spin column(s), followed by centrifugation at 11,000 g for 30 seconds, after which the flow-through was discarded. An incubation mix was prepared in a 1.7 mL Eppendorf tube by combining 0.01 mL of DNase I stock solution with 0.07 mL of Buffer RDD. Next, 0.08 mL of the incubation mix was added to the spin column membrane, and the columns were incubated at RT for 15 minutes. Following incubation, 0.35 mL of RW1 buffer was added to the spin column(s), and the columns were centrifuged at 11,000 g for 30 seconds, with the flow-through discarded. Then, 0.50 mL of RPE buffer was added to the spin column(s) and centrifuged at 11,000 g for 30 seconds, and the flow-through was discarded. This step was repeated. The spin column(s) were then transferred to a new 2 mL collection tube and centrifuged at maximum speed for 1 minute at 4°C. The spin column(s) were subsequently placed into tubes, and RNA was eluted by adding 0.02 mL of nuclease-free water followed by centrifugation at 11,000 g for 1 minute. cDNA synthesis was performed using SMART® MMLV reverse transcriptase (Takara, 639524) with oligo(dT) primers. Quantitative real-time PCR was conducted on a ViiA 7 Real-Time PCR System (Life TechnologiesTM - Applied Biosystems®) using Power SYBR™ Green PCR Master Mix (Life Technologies, 4367660). Transcript levels were normalized to Rpl13a expression (housekeeping). Primer sequences used are provided in Primers Table.

### Sennoside A in vivo treatment

#### Acute

Sennoside A (SA, Sigma-Aldrich, 68909) was initially reconstituted in dimethyl sulfoxide (DMSO). SA was administered intragastrically (i.g) at 10 mg/kg or 30 mg/kg in drinking water, in a final volume of 150 µL. Control animals received the same volume of the vehicle solution alone (drinking water + DMSO). Oral gavage was performed using plastic needles (Fine Science Tools, 18961-20) during the dark phase, at zeitgeber time 14-15 (ZT14-15).

#### Chronic

Mice assigned to SA treatment received 20 mg/kg SA in a final volume of 150 µL by intragastric gavage 24 h before each oral OVA challenge. SA was administered on days 13, 16, 20, 23, and 27. The corresponding control groups received an equivalent volume of vehicle containing DMSO diluted in water.

### Allergic sensitization and treatment

5-6-week-old female BALB/c mice were sensitized subcutaneously on days 0 and 7 with endotoxin-free ovalbumin (OVA; 0.25 mg/kg; BioVendor, catalog no. 321001) adsorbed to aluminum hydroxide gel (alum; 50 mg/kg; InvivoGen, catalog no. vac-alu-250). The sensitization solution was diluted in phosphate-buffered saline (PBS; pH 7.4) to a final administration volume of 200 µL per mouse. Control mice received alum diluted in PBS without OVA and are referred to as the PBS/alum group.

All mice were orally challenged by intragastric gavage with 40 mg of grade III OVA (Sigma-Aldrich, catalog no. A5378) dissolved in water to a final volume of 150 µL. Oral challenges were performed on days 14, 17, 21, 24, and 28. Allergic sensitizations and oral allergen challenges were performed at zeitgeber time 16 (ZT16), corresponding to 4 h after the beginning of the dark phase.

### Temperature

Temperature was measured using a rectal probe (Digi-Sense Model WD 20250-91). For acute sennoside A exposure, mice were monitored every 30 minutes for up to 6 h post-gavage. For the OVA model, rectal temperature was measured immediately following the fifth oral OVA challenge and every 15 minutes for 2 h post-gavage. A water-based lubricant (Lubricant Liquid, CVS Health) was applied to the probe to minimize irritation, and the probe was sanitized with 70% ethanol between animals.

### Diarrhea

Diarrhea was determined by visual assessment of stool consistency, wetness, and frequency upon oral exposure to sennoside A and vehicle solution. Diarrhea were unbiasedly determined by a team member unaware of the cohort identities.

### Tissue RNA extraction and quantification

Tissues were harvested in RNA STAT-60 RNA isolation reagent (Amsbio, CS-502-500) and homogenized by ceramic bead homogenization (Omni, INC). RNA was extracted using Direct-zol RNA Mini Prep (ZYMO Research, R2072). cDNA synthesis was performed using SMART® MMLV reverse transcriptase (Takara, 639524) with oligo(dT) primers. Quantitative real-time PCR was conducted on a ViiA 7 Real-Time PCR System (Life TechnologiesTM - Applied Biosystems®) using Power SYBR™ Green PCR Master Mix (Life Technologies, 4367660). Transcript levels were normalized to Rpl13a expression (housekeeping).

### Blood and tissue collection

Mice were euthanized 24 h after the final oral OVA challenge on day 29. Terminal blood was collected via cardiac puncture and allowed to clot before serum was isolated by centrifugation. Serum samples were stored at -80C until analysis.

The duodenum and jejunum were collected immediately after euthanasia, homogenized in RNAZol (RN 190 MCR) for subsequent gene-expression analysis.

### Serum antibody measurements

Serum was collected by cardiac puncture 24 h after the fifth oral OVA gavage and assayed for total IgE, OVA-specific IgE, and OVA-specific IgG1 by enzyme-linked immunosorbent assay (ELISA). For total IgE and OVA-specific IgE, ELISA-grade plates (MaxiSorp Nunc, Thermo Scientific) were coated with 2 μg/mL purified rat anti-mouse IgE antibody (clone R35-72; BD Pharmingen, 553413) in 0.1 M sodium carbonate buffer (pH 9.5) overnight at 4°C. Plates were blocked with PBS containing 1% heat-inactivated fetal bovine serum for 2 h at room temperature (RT). Serum was diluted 1:10 or 1:50 and incubated for 2 h at RT. For total IgE, purified mouse IgE (BD Biosciences, 557080) was used to generate a standard curve, with the highest concentration being 500 ng/mL followed by two-fold serial dilutions. An anti-OVA IgE monoclonal antibody (Bio-Rad, MCA2259) was used to generate a standard curve, with the highest concentration being 1,000 ng/mL followed by two-fold serial dilutions. Total IgE was detected with 0.5 μg/mL biotinylated rat anti-mouse IgE antibody (clone R35-118; BD Pharmingen, 553414) for 1 h at RT. OVA-specific IgE was detected with biotinylated OVA (OVAI-BU-1), diluted 1:250 from a 2 mg/mL stock, for 1 h at RT.

For OVA-specific IgG1, plates were coated with 20 μg/mL grade III OVA (Sigma-Aldrich, A5378) in coating buffer overnight at 4°C. Standard wells were coated with purified mouse IgG1 monoclonal antibody (BD Biosciences, 557273), with the highest concentration being 500 ng/mL followed by two-fold serial dilutions. Plates were blocked with assay diluent for 1 h at RT. Serum was diluted 1:10,000 and incubated for 1 h at RT. OVA-specific IgG1 was detected with biotinylated rat anti-mouse IgG1 antibody (clone A85-1; BD Biosciences, 553441) diluted 1:5,000 for 1 h at RT.

For all assays, HRP-conjugated streptavidin (1:1,000; BD Biosciences, 554066) was incubated for 30 min at RT. Plates were then developed with TMB substrate reagent (BD Biosciences, 555214) in the dark at RT, with color development checked every 1 min for up to 15 mins. Reactions were stopped with 2.5 N sulfuric acid (Ricca, UN2796) at the appearance of a two-fold dilution pattern on the standard curve. Absorbance at 450 nm was read immediately using a Varioskan LUX plate reader (Thermo Scientific, Waltham, MA). Between each step, plates were washed three to seven times with 0.05% Tween-20 (Sigma-Aldrich, P1379) in PBS. Concentrations were calculated from the standard curve generated for each assay.

### Primers Table

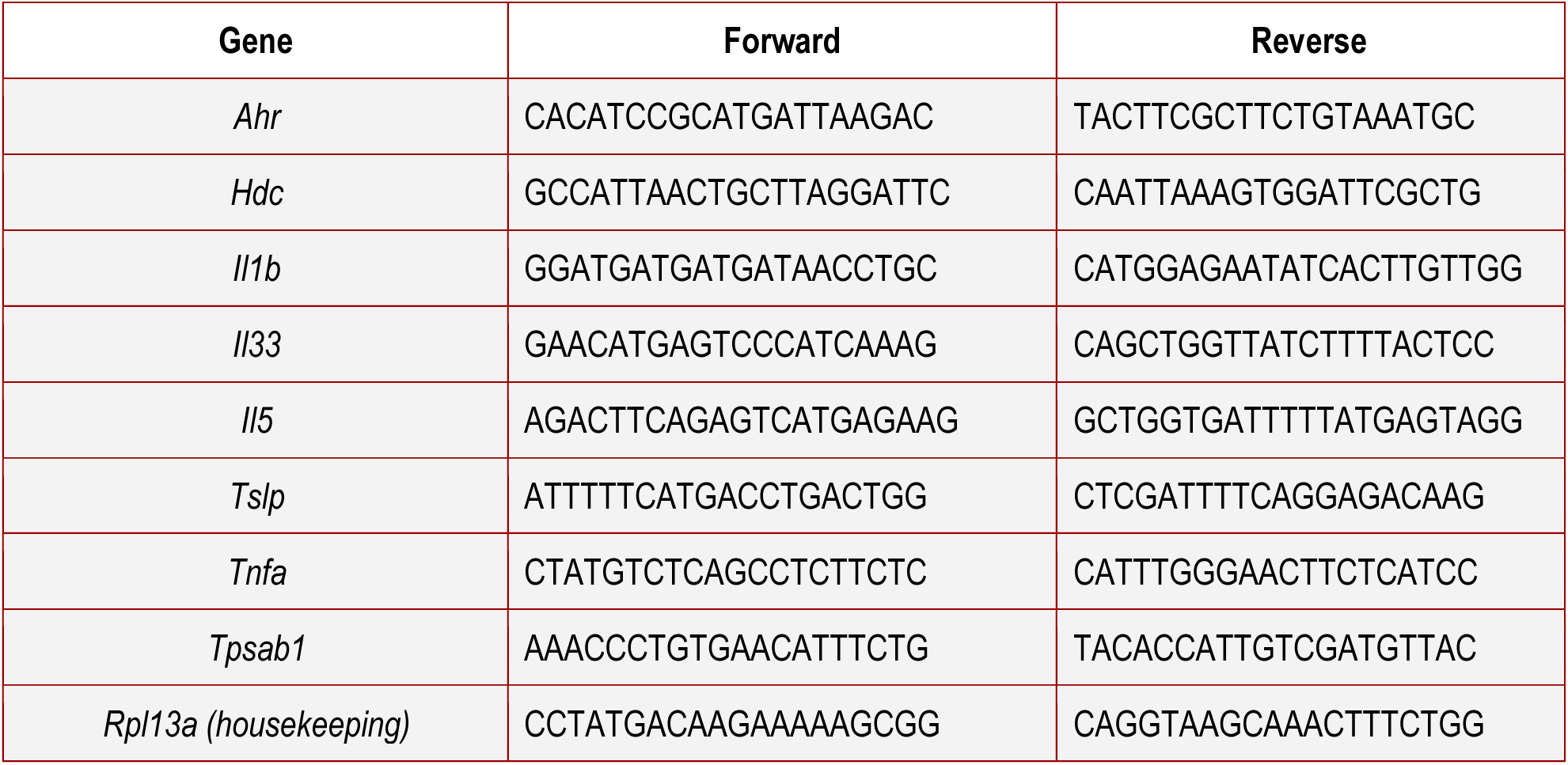

### Statistical analysis

Statistical analyses were carried out in GraphPad Prism software v10.1.0. Data were analysed with the Mann–Whitney U-test. Statistical significance is defined as *p < 0.05, **p < 0.01, ***p < 0.001, and ****p < 0.0001. Nonparametric statistical analyses were used throughout the manuscript, and all data are mean ± s.e.m., unless stated otherwise.

## ACKNOWLEDGMENTS

We would like to thank all current and former members of the Florsheim lab for helpful discussions. We thank the Department of Animal Care and Technologies team for technical assistance, and Biodesign Institute employees for their continuous assistance. F.N. received support from the School of Life Sciences Undergraduate Research from Arizona State University. Work in the Florsheim lab was supported by the Food Allergy Science Initiative (2022-FASI-0001, 2023-FASI-AS23310), the Hypothesis Fund (G11391-300), and the Arizona Department of Health (2024-022). Schematics were created with BioRender.com.

## Author contributions

F.N. and E.B.F. designed the study, analyzed the data, and wrote the manuscript with input from the other authors. F.N. performed *in vitro* experiments. A.C.R. performed *in vivo* acute sennoside A experiments. C.W. performed *in vivo* gut allergic inflammation experiments. E.B.F. supervised the research.

## Declaration of generative AI and AI-assisted technologies

During preparation of this manuscript, the authors used ChatGPT (OpenAI) to improve conciseness, grammar, and readability. All suggested revisions were reviewed and edited by the authors, who take full responsibility for the final content of the manuscript.

## SUPPLEMENTAL FIGURE LEGENDS

**Supplementary Figure 1 | Flow cytometry characterization of BMMCs** Flow cytometry was used to characterize the phenotype of bone marrow–derived mast cells (BMMCs) generated in vitro. Cells were stained with PE conjugated anti-FcεR1 and APC-Cy7 conjugated anti-cKit (CD117). Cells were gated to identify single cells, then single-color controls using stained cells were established for positive detection of CD117 and FcεR1. Note that greater than 90% of BMMCs are FcεR1+ cKit+.

