## Supplemental Figure 1 for "Sennoside A differentially regulates mast cell activation and attenuates gut allergic inflammation"

### Supplementary Figure 1

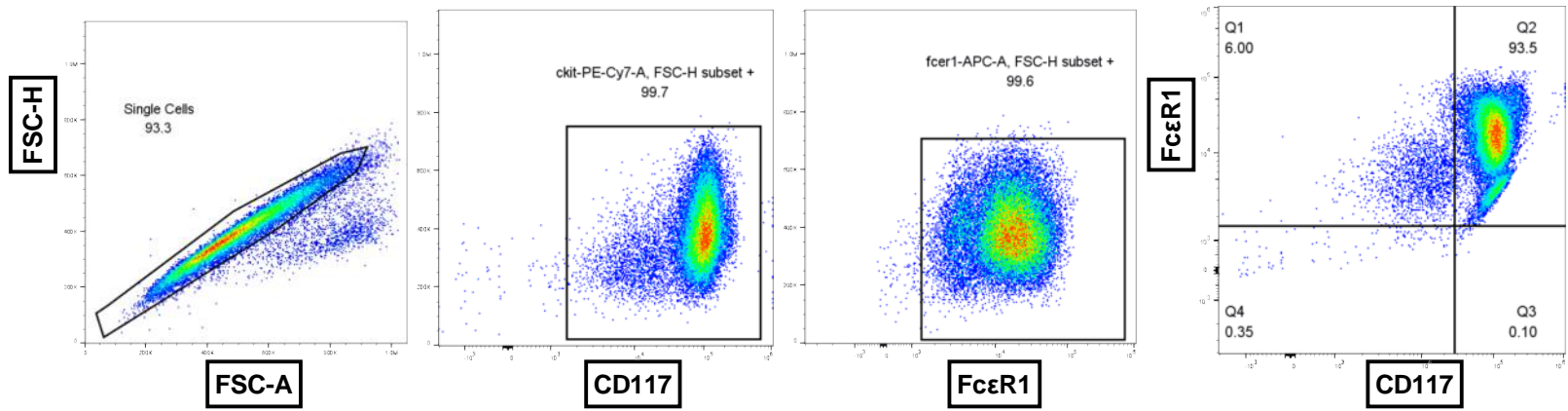

#### Supplementary Figure 1 | Flow cytometry characterization of BMMCs

Flow cytometry was used to characterize the phenotype of bone marrow-derived mast cells (BMMCs) generated in vitro. Cells were stained with PE conjugated anti-FcεR1 and APC-Cy7 conjugated anti-cKit (CD117). Cells were gated to identify single cells, then single-color controls using stained cells were established for positive detection of CD117 and FcεR1. Note that greater than 90% of BMMCs are FcεR1+ cKit+
